# Δ^9^-tetrahydrocannabinol Attenuates Oxycodone Self-Administration Under Extended Access Conditions

**DOI:** 10.1101/239038

**Authors:** Jacques D. Nguyen, Yanabel Grant, Kevin M. Creehan, Candy S. Hwang, Sophia A. Vandewater, Kim D. Janda, Maury Cole, Michael A. Taffe

## Abstract

Growing nonmedical use of prescription opioids is a global problem, motivating research on ways to reduce use and combat addiction. Medical cannabis (“medical marijuana”) legalization has been associated epidemiologically with reduced opioid harms and cannabinoids have been shown to modulate effects of opioids in animal models. This study was conducted to determine if Δ^9^-tetrahydrocannabinol (THC) enhances the behavioral effects of oxycodone.

Male rats were trained to intravenously self-administer (IVSA) oxycodone (0.15 mg/kg/infusion) during 1 h, 4 h or 8 h sessions. Following acquisition rats were exposed to THC by vapor inhalation (1 h and 8 h groups) or injection (0-10 mg/kg, i.p.; all groups) prior to IVSA sessions. Fewer oxycodone infusions were obtained by rats following vaporized or injected THC compared with vehicle treatment prior to the session. Follow-up studies demonstrated parallel dose-dependent effects of THC, i.p., on self-administration of different per-infusion doses of oxycodone and a preserved loading dose early in the session. These patterns are inconsistent with behavioral suppression. Additional groups of male and female Wistar rats were assessed for nociception following inhalation of vaporized THC (50 mg/mL), oxycodone (100 mg/mL) or the combination. Tail withdrawal latency was increased more by the THC/oxycodone combination compared to either drug alone. Similar additive antinociceptive effects were produced by injection of THC (5.0 mg/kg, i.p.) and oxycodone (2.0 mg/kg, s.c.). Together these data demonstrate additive effects of THC and oxycodone and suggest the potential use of THC to enhance therapeutic efficacy, and to reduce the abuse, of opioids.

## 1. Introduction

Nonmedical opioid abuse is a significant global problem, with an estimated 33 million users of opiates and prescription opioids worldwide (UNODC, 2016). Approximately 2 million people in the US have a prescription opioid related abuse disorder (CBHSQ, 2015), which may increase the likelihood of later non-prescription opioid use (Muhuri et al., 2013), and prescription opioid related overdose deaths have drastically increased over the last two decades (CDC, 2016). Despite the growing impact of prescription opioids on public health, relatively few pre-clinical studies have investigated the self-administration of oxycodone, one of the most commonly prescribed medications (OxyContin® or as part of Percocet®). Available studies confirm that oxycodone self-administration causes reward-related behavioral changes (Zhang et al., 2016) sometimes physical dependence and withdrawal (Enga et al., 2016) in mice and that male and female rats acquire oxycodone self-administration at similar rates (Mavrikaki et al., 2017). Thus, traditional self-administration models can be used to evaluate approaches to reduce prescription opioid abuse.

Indirect evidence suggests that cannabis (“marijuana”) may attenuate some of the harms associated with opioid use. Epidemiological studies have recently reported reductions in opioid positive drivers in car crash fatalities in younger drivers 21-40 (Kim et al., 2016) over nonmedical marijuana states and in-patient hospitalization rates for opioid dependence were 23% lower in medical marijuana states compared with nonmedical marijuana states (Shi, 2017). Opioid overdose mortality is lower in states with medical marijuana legalization (Bachhuber et al., 2014) and an experimental study found that inhalation of cannabis (via Volcano® vaporizer) decreased pain in chronic pain patients that were being maintained on extended release oxycodone or morphine without changing the plasma concentration-time curves for either medication (Abrams et al., 2011). A recent clinical study found that smoked cannabis enhanced the analgesic effects of oxycodone and produced modest increases in positive subjective ratings related to oxycodone when administered in combination (Cooper et al., 2018). These findings suggest that psychoactive cannabinoids may interact with the effects of opioids, both to enhance therapeutic impact and to potentially reduce nonmedical use.

Currently there is only limited direct evidence for the interactive effects of cannabinoid and opioid receptor signaling, as reviewed in (Scavone et al., 2013; Wenzel and Cheer, 2018); a few preclinical studies have investigated whether cannabinoid receptor activation via CB1 agonists, including Δ^9^-tetrahydrocannabinol (THC), can modify the effects of heroin or morphine but no studies have investigated the combined effects of THC with oxycodone. Daily treatment with THC by injection decreased responding for intravenous heroin in rhesus monkeys (Maguire and France, 2016) and similar effects were observed after injection of full agonist CB1 ligands (Maguire et al., 2013). Solinas and colleagues have shown that THC (3 mg/kg, i.p.) decreases heroin intravenous self-administration (IVSA) under a fixed-ratio 1 (FR1) schedule; however, they have also observed facilitation of heroin IVSA under progressive ratio (PR) schedule of reinforcement and following specific experimental approaches such as limited training sessions (10 days of acquisition; short access 3 h) and food restriction conditions (Solinas and Goldberg, 2005; Solinas et al., 2004). In addition, THC enhanced the antinociceptive effect of morphine in mice (Pugh et al., 1996), and the cannabinoid receptor full-agonists CP55,940 and WIN 55,212 enhanced antinociceptive effects of morphine in rhesus monkeys (Li et al., 2008; Maguire and France, 2014). Inhibitors of endocannabinoid catabolic enzymes may attenuate heroin and morphine-induced antinociception and dependence in mice (Ramesh et al., 2013; Wilkerson et al., 2017). This predicts that THC may likewise interact with the effects of prescription opioids such as oxycodone.

This study was therefore designed to determine if THC inhalation increases oxycodone-induced antinociception and alters oxycodone IVSA. Because THC is typically administered via inhalation in humans, and by parenteral injection in rat models, it is of significant interest to determine if the two routes of administration produce similar interactions of THC with oxycodone. A new method for delivery of drugs to rats using electronic cigarette (e-cigarette) technology has been recently reported (Nguyen et al., 2016a; Nguyen et al., 2016b) and, pursuant to this study, inhaled THC produced antinociceptive effects commensurate with those produced by 10 mg/kg THC, i.p. (Javadi-Paydar et al., 2018; Nguyen et al., 2016b). Male and female rats were evaluated on a tail withdrawal response assay to determine any interactive effects of THC and oxycodone on nociception. To determine potential interactive effects on reward we used multiple IVSA models. First, we used an extended-access paradigm in which rats were permitted to self-administer oxycodone in 8 h sessions as a stronger test of effects of THC on a compulsive-like behavioral phenotype (Vendruscolo et al., 2011; Wade et al., 2015). Additional studies were completed in 1 h and 4 h oxycodone access groups to determine if any observed interactions were most likely due to acute pharmacodynamical interaction or if they required, e.g., changes associated with the escalated phenotype. The study further sought to confirm receptor specificity by evaluating the effect of pre-treatment with the mu opioid receptor antagonist, naloxone, non-contingent doses of oxycodone and the cannabinoid receptor 1 (CB1) inverse agonist/antagonist, SR141716 (Rimonabant). Repeated vapor inhalation of THC during adolescence induces a lasting tolerance to THC in adulthood (Nguyen et al., 2018a; Nguyen et al., 2018b) and repeated THC injection during adolescence increases heroin IVSA as adults (Ellgren et al., 2007). Studies were therefore conducted in a group of adult rats that were exposed to THC or PG vapor in adolescence (Nguyen et al., 2018a), but self-administered approximately equal amounts of oxycodone to determine any differential effect of THC. Several strategies were used to address the potential concern that general sedative effects of THC would obscure any interaction with oxycodone in the IVSA model. First, we chose vapor inhalation regimens (Javadi-Paydar et al., 2018; Nguyen et al., 2016b) and injection doses (Taffe et al., 2015; Wiley and Burston, 2014; Wiley et al., 2007) that have been shown not to significantly alter locomotor behavior in rats. Examination of the behavioral patterns generated early in the IVSA sessions was used to further dissociate an inability to respond on the lever from a reduction in oxycodone rewards obtained. We also used two strategies that reduce the oxycodone penetration to the brain for each response, anti-oxycodone vaccination (Nguyen et al., 2018c) and per-infusion dose reduction, thereby requiring increased behavioral output; parallel effects of THC on IVSA would be incompatible with an inference of general sedation, if observed.

## 2. Methods

### 2.1 Subjects

Male (N=76) and female (N=8) Wistar (Charles River, New York) rats were housed in humidity and temperature-controlled (23±1 °C) vivaria on 12:12 hour light:dark cycles. Animals entered the laboratory at 10-11 weeks of age except for the N=24 males that arrived on post-natal day (PND) 22. Animals had ad libitum access to food and water in their home cages and all experiments were performed in the rats’ scotophase. All procedures were conducted under protocols approved by the Institutional Care and Use Committees of The Scripps Research Institute and in a manner consistent with the Guide for the Care and Use of Laboratory Animals (National Research Council (U.S.). Committee for the Update of the Guide for the Care and Use of Laboratory Animals. et al., 2011).

### 2.2 Drugs

(-)-Oxycodone HCl and (-)-naloxone HCl were obtained from Sigma-Aldrich (St. Louis, MO). SR141716 was purchased from Cayman Chemical. The Δ^9^-tetrahydrocannabinol (THC) and heroin HCl were obtained from NIDA Drug Supply. For injection experiments, THC was supplied in ethanolic solution (50-200 mg/mL); the ethanol was evaporated off prior to preparation in a 1:1:8 ratio of ethanol:cremophor:saline. Oxycodone, heroin and naloxone were dissolved in saline (0.9%), for injection. For vapor inhalation experiments, drugs were dissolved in propylene glycol (PG) with the concentrations expressed as mg of drug per mL of PG. PG was obtained from Sigma-Aldrich (St. Louis, MO). Drug injections were administered, and vapor inhalation sessions were initiated, 30 min prior to the start of self-administration sessions.

### 2.3 Inhalation Apparatus and Procedure

The vapor inhalation was conducted using an approach which has been previously described (Javadi-Paydar et al., 2019; Javadi-Paydar et al., 2018; Nguyen et al., 2016a; Nguyen et al., 2016b; Nguyen et al., 2018b). In brief, vacuum controlled exhaust (i.e., a “pull” system) flowed room ambient air through a sealed rat chamber. A stream of vapor derived from an e-cigarette atomizer/cartridge was then integrated with the ambient air stream once triggered by computerized control using a proprietary interface module (La Jolla Alcohol Research, Inc, La Jolla, CA, USA). Rats were exposed to 30 min of THC (12.5-200 mg/mL in the propylene glycol vehicle) vapor inhalation (followed by a 5 min period for chamber clearance) immediately prior to the start of self-administration sessions or the first antinociception assessment. Additional methodological detail is provided in the **Supplemental Methods**.

### 2.4 Repeated Adolescent Vapor Exposure

Adolescent rats (N=12 per group) were exposed to twice daily vapor inhalation sessions (30 minutes; THC (100 mg/mL) or the propylene glycol (PG) vehicle) on post-natal days (PND) 35-39 and PND 42-46, as is described in detail in the **Supplemental Methods**. These groups initiated IVSA of oxycodone (0.15 mg/kg/infusion; 8 h sessions; Fixed Ratio 1 response contingency) on PND 112 for a study described in a preprint (Nguyen et al., 2018a). Following acquisition and initial dose substitution studies (see **Supplemental Methods**) the rats were returned to 4 h sessions of oxycodone IVSA (0.15 mg/kg/infusion; FR1) for the current study because results of the 8 h IVSA group showed that effects of THC pre-treatment only lasted for about 4-5 hours. Rats were pre-treated with doses of THC (0.0, 1.0, 5.0, 7.5, 10.0 mg/kg, i.p.), 30 minutes prior to the start of the self-administration sessions in a counter-balanced order. Next, rats were allowed to self-administer oxycodone at a lower dose (0.06 mg/kg/infusion; FR1; 4 h sessions) and the THC pre-treatment dose series completed again. A total of 19 rats (N=9, repeated THC group) completed the study with patent catheters.

### 2.5 Anti-oxycodone vaccination

Rats were administered a conjugate vaccine (Oxy-TT; N=12) or the tetanus toxoid carrier protein only (TT; N=10) on Weeks 0, 2, and 4, adapted from a vaccination protocol previously reported (Nguyen et al., 2018c); see **Supplemental Methods** for full details. Rats were prepared with chronic intravenous catheters on week 7 of the vaccination protocol and allowed one week of recovery prior to the start of self-administration experiments. Within the TT group, 8 rats completed acquisition with patent catheters and 7 completed the THC inhalation study. Within the Oxy-TT group, 11 completed the entire study.

### 2.6 Self-administration procedure

Rats were prepared with chronic indwelling intravenous catheters using sterile procedures under gas anesthesia as described previously (Nguyen et al., 2017) and in the **Supplemental Methods.** Briefly, the catheter tubing was passed subcutaneously from a port at the back and inserted into the jugular vein. *A minimum of 4 days was allowed for surgical recovery prior to starting an experiment.* Catheters were flushed with ~0.2-0.3 ml heparinized (166.7 USP/ml) saline before sessions and ~0.2-0.3 ml heparinized saline containing cefazolin (100 mg/mL) after sessions. Catheter patency was assessed once a week after the last session of the week, via administration of the ultra-short-acting barbiturate anesthetic Brevital sodium (1% methohexital sodium; Eli Lilly, Indianapolis, IN). Drug self-administration was conducted in operant boxes (Med Associates) located inside sound-attenuating chambers located in an experimental room (ambient temperature 22 ± 1 °C; illuminated by red light) outside of the housing vivarium. To begin a session, the catheter fittings on the animals’ backs were connected to polyethylene tubing contained inside a protective spring suspended into the operant chamber from a liquid swivel attached to a balance arm. Each operant session started with the extension of two retractable levers into the chamber. Following each completion of the response requirement (response ratio), a white stimulus light (located above the reinforced lever) signaled delivery of the reinforcer and remained on during a 20-sec post-infusion timeout, during which responses were recorded but had no scheduled consequences. Drug infusions were delivered via syringe pump. The training dose (0.15 mg/kg/infusion; ~0.1 ml/infusion) was selected from prior self-administration studies (Nguyen et al., 2018c; Wade et al., 2015). The first group of naïve male rats (N=12), and the adolescent vapor-exposure groups (N=12 per group) of male rats were trained in 8 h sessions using a Fixed Ratio 1 response contingency. The tetanus toxoid (TT) control (N=10) and oxycodone-TT (N=12) vaccinated groups of male rats were trained in 1 h sessions under a Progressive Ratio (PR) response contingency for the initial 7 sessions and Fixed Ratio 1 thereafter. In the PR paradigm, the required response ratio was increased after each reinforcer delivery within a session (Hodos, 1961; Segal and Mandell, 1974) as determined by the following equation (rounded to the nearest integer): Response Ratio=5e^(injection number\**j*)–5 (Richardson and Roberts, 1996). The *j* value was set to 0.2. This group was initially trained with a PR contingency since a prior study (Nguyen et al., 2018c) had found that the Oxy-TT vaccination resulted in higher IVSA oxycodone self-administration under FR response contingency but a greater reduction in the amount of drug self-administered associated with the increased workload of a PR paradigm. Rats were trained during weekdays (5 days per week).

### 2.7 Nociception Assay

Tail withdrawal antinociception was assessed using a water bath (Bransonic® CPXH Ultrasonic Baths, Danbury, CT) maintained at 52 °C. The latency to withdraw the tail was measured using a stopwatch and a cutoff of 15 seconds was used (Wakley and Craft, 2011; Wakley et al., 2014). Tail withdrawal was assessed starting 35 minutes after the initiation of inhalation or 30 minutes after injection. Nociception experiments following injected oxycodone (1,2 mg/kg, s.c.), THC (5,10 mg/kg, i.p.) or the combination were conducted in a group of adult male (N=6; 48 wks of age, 665.2, SD 76.2 g) and female (N=8; 48 wks of age, 310.5, SD 44.3 g) Wistar rats that were previously used in experiments of chronic vapor inhalation of THC (Nguyen et al., 2018b). Nociception experiments following vapor inhalation were conducted in a separate group of male Wistar rats (N=10; 44 wks of age, 722.6, SD 84.2 g), that were previously used in pilot experiments with the nociception assay following vapor inhalation of heroin, oxycodone, methadone and THC to determine exposure conditions.

### 2.8 Pharmacokinetics

A separate group (N=6) male rats were prepared with indwelling intravenous catheters for serial blood sampling. During experiments, blood samples (~250 µl) were obtained at four time points (35, 60, 120 and 240 minutes) after injection or the start of vapor inhalation. Experiments were conducted within-group with a minimum 14 day interval between them to accord with recommendations for blood volume recovery (Diehl et al., 2001). Plasma levels of THC were assessed by LCMS as previously described (Javadi-Paydar et al., 2018; Nguyen et al., 2016b) and as detailed in the **Supplemental Methods**.

### 2.9 Data Analysis

Analysis of IVSA data was conducted with Analysis of Variance (ANOVA) on the number of infusions obtained. Within-subjects factors of Session and Drug Treatment condition were included as relevant. Between-groups factors of vaccine treatment and adolescent vapor inhalation condition were included for their respective studies. A Grubbs test eliminated two individuals from the vaccine study which exhibited outlier IVSA during the vapor-inhalation test (Drug infusions 2.9 SD greater than the mean were observed for one individual on the Air and one on the THC condition). Tail withdrawal latencies and plasma THC levels were analyzed with repeated measures ANOVA including within-subjects factors of Drug Treatment Condition (or Route of Administration) and Time post-injection/initiation of vapor. Significant main effects were followed with post hoc analysis using Dunnett, Tukey (multi-level factors) or Sidak (two-level factors) tests for multiple comparisons. All analysis used Prism 6 or 7 for Windows (v. 6.07 and 7.04; GraphPad Software, Inc, San Diego CA).

## 3. Results

### 3.1 THC vapor attenuates oxycodone self-administration under extended access (8 h) conditions

Male rats (N=11) trained to self-administer oxycodone in 8 h sessions significantly escalated their self-administration during 17 sessions of acquisition training [F(3.255,32.55)=11.41; *p*<0.0001] as is shown in **Figure 1A.** The post hoc analysis confirmed significant increases in oxycodone self-administration relative to the first session across sessions 6-17. Inhalation of THC (200 mg/mL; 30 min) immediately prior to the self-administration session significantly reduced the mean (N=9; two rats were excluded due to mechanical failure on one of the test sessions) infusions of oxycodone compared to the effect of inhalation of the PG vehicle (**Figure 1B**). The ANOVA confirmed significant effect of vapor Treatment [F(1,8)=10.27; *p*<0.05] and of the interaction of factors [F(7,56)=5.587; *p*<0.001]. Post hoc analysis confirmed that oxycodone IVSA was reduced in hours 1-3 and 5 relative to the PG inhalation condition. There was no difference in lever discrimination observed between PG and THC-exposed rats (76.59±3.85 and 73.92±7.33 percent, respectively). Furthermore, a second study showed that inhalation of vaporized THC (100 and 200 mg/ml) significantly attenuated oxycodone self-administration (**Figure 1C**) compared to inhalation of PG vehicle and in a dose-dependent manner. In this study the vapor administration schedule was randomized for PG and 200 mg/mL on T, Th, with 100 mg/ml for all rats on the next F. Inhalation of 12.5 and 25 mg/ml THC (not shown) did not significantly alter oxycodone self-administration. The rmANOVA confirmed significant main effects of Time [F(7,70)=2.501; *p*<0.05], of vapor Treatment [F(2,20)=6.016; *p*<0.01] and of the interaction of factors [F(14,140)=2.322; *p*<0.01]. Similarly, pre-session injection of THC (0, 5, 10 mg/kg, i.p.) significantly reduced oxycodone self-administration (**Figure 1D**). The analysis confirmed significant effects of Time [F(7,21)=2.729; *p*<0.01], of vapor Treatment [F(2,30)=7.168; *p*<0.01] and of the Time x Treatment interaction [F(30,210)=7.142; *p*<0.0001]. The post hoc analysis further confirmed that oxycodone IVSA was significantly reduced for up to 5 h in rats pretreated with THC 10 mg/kg., i.p..

**Figure 1.**
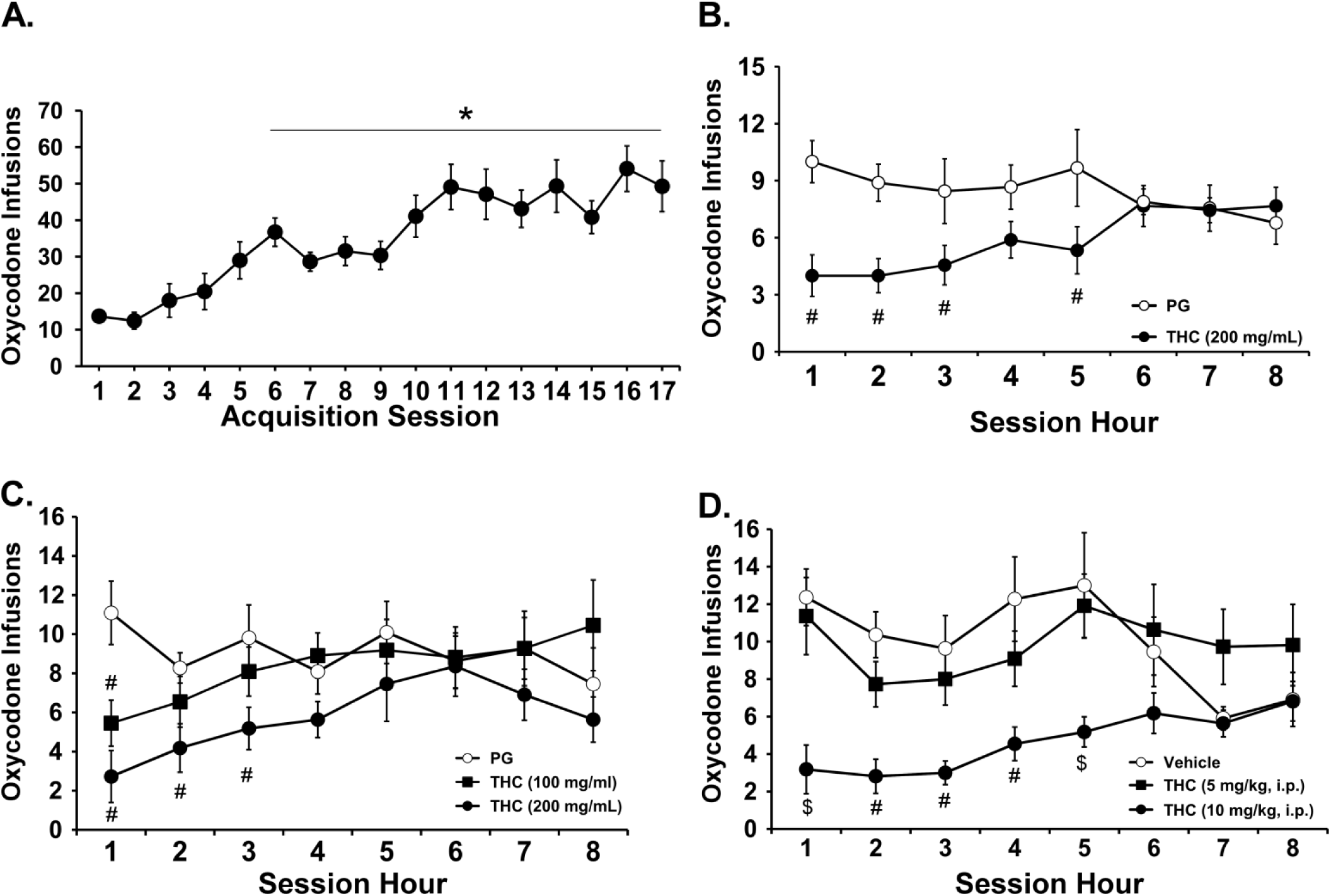
THC reduces oxycodone self-administration under extended access conditions. A) Mean (N=11; ±SEM) infusions for male rats trained to self-administer oxycodone (0.15 mg/kg/inf) within 8 h extended access. B) Mean (N=11; ±SEM) infusions following THC vapor. Significant differences within group from session 1 are indicated by *. Significant differences from PG vehicle condition are indicated by *. Mean (N=11; ±SEM) infusions for male rats trained to self-administer oxycodone following C) THC vapor (100,200 mg/ml) and D) injected THC (5,10 mg/kg, i.p.). Significant differences from vehicle condition are indicated by # and from all other dose conditions with $.

### 3.2 THC vapor attenuates oxycodone self-administration via CB1 receptor activation

The first group continued with 8 h self-administration sessions for the evaluation of the effects of the CB1 receptor antagonist, SR141716 (SR), on the THC-induced reduction of oxycodone IVSA. Analysis focused on the first 90 minutes of the session because of the anticipated duration of action of the SR and indeed there were no apparent effects of SR beyond this interval. The data show that the effects of THC were attenuated by systemic administration of SR (4 mg/kg, i.p.) prior to the vapor inhalation session (**Figure 2**). The ANOVA confirmed a significant main effect of Treatment [F(2,20)=7.829; *p*<0.01] and post hoc analysis of the marginal means confirmed that oxycodone self-administration following veh-THC pre-treatment was significantly lower than after either SR-PG or SR-THC pre-treatment conditions.

**Figure 2.**
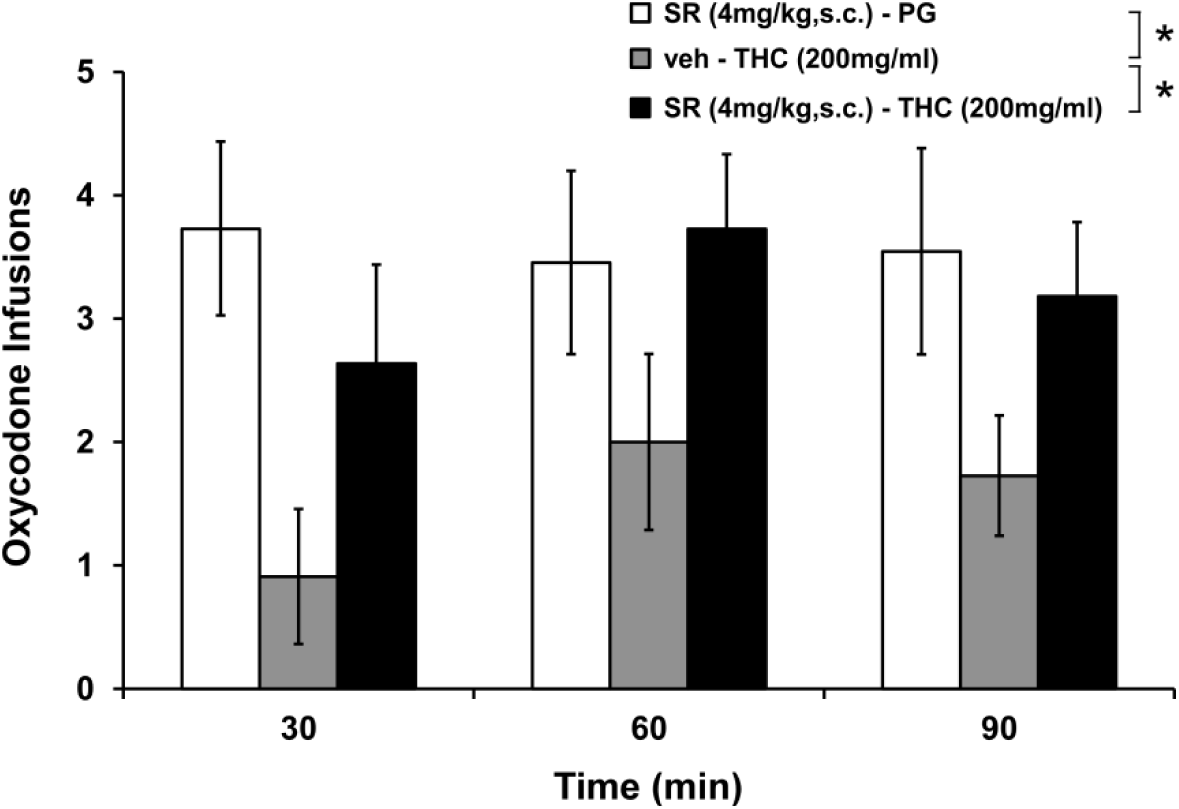
THC-mediated attenuation of oxycodone self-administration is CB_1_ receptor-mediated. Mean (N=11; ±SEM) infusions of oxycodone following vapor inhalation of THC and injection of CB1 antagonist, SR141716 (SR; 4 mg/kg, i.p.) prior to the inhalation session. A significant difference between treatment conditions (across time bins) is indicated with *.

### 3.3 Mu opioid receptor agonism and antagonism

Experiments were conducted to determine the effects of pre-treating animals with mu opioid receptor (MOR) agonist (oxycodone) or MOR antagonist (naloxone) compounds. Pretreatment with oxycodone (0, 0.5, 1, 2 mg/kg, i.p.) significantly attenuated oxycodone IVSA in a dose-dependent manner (**Figure 3A**) and statistical analysis of the first 2 h confirmed a significant effect of oxycodone Treatment [F(3,18)=8.401; *p*<0.01]. Post hoc analysis of the marginal means confirmed that oxycodone self-administration after the highest pre-treatment dose (2.0 mg/kg, s.c.) was significantly lower than all other conditions.

**Figure 3.**
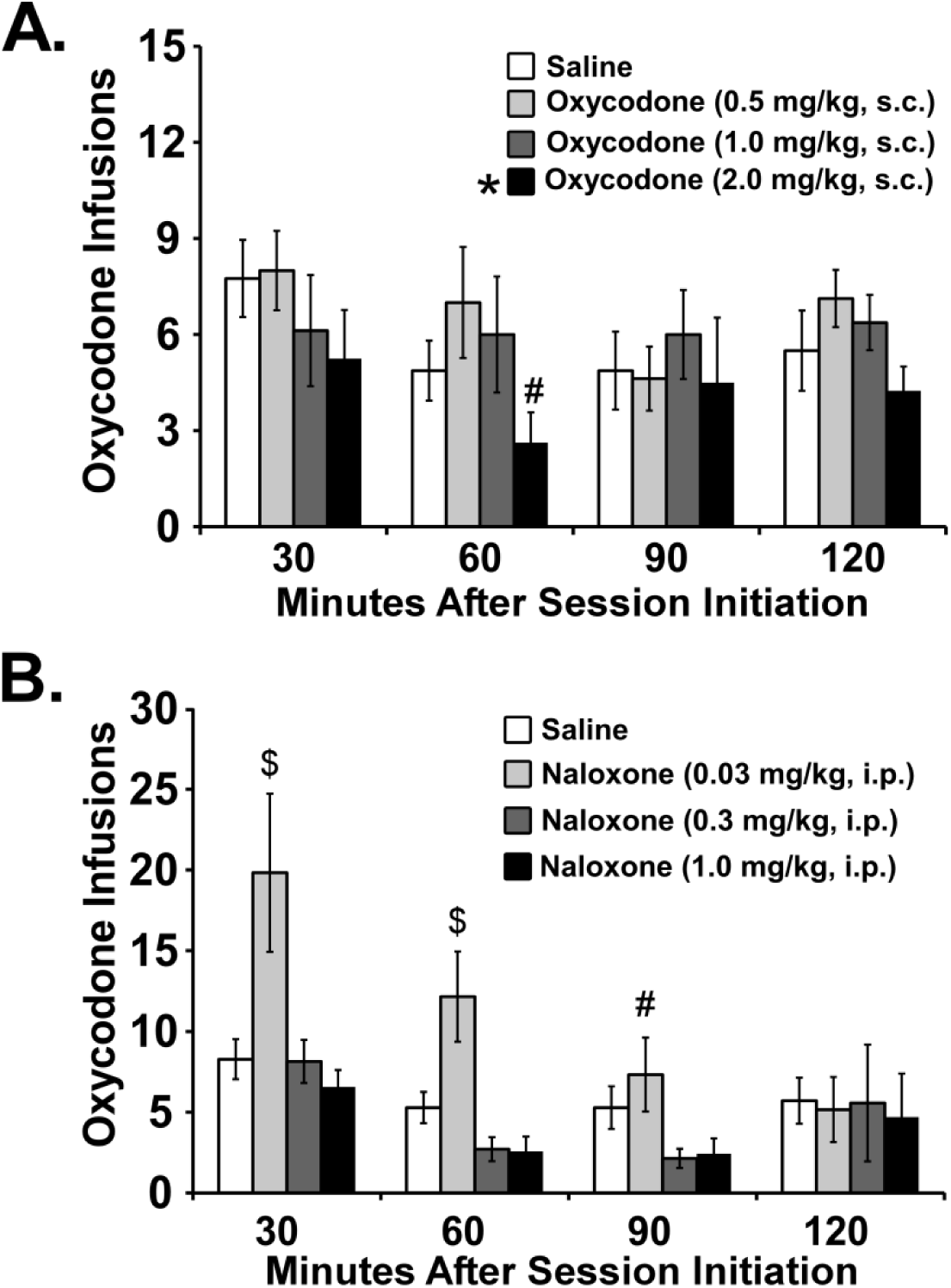
Mu opioid receptor agonists attenuate oxycodone self-administration. Mean (±SEM) infusions of oxycodone following injection of A) oxycodone (N=8; 0.5-2 mg/kg, s.c.) or B) naloxone (N=6; 0.03-1.0 mg/kg, i.p.). A significant difference from the first hour time point is indicated by *. A significant difference from all other dose conditions is indicated with $ and a difference from the 0.3 is indicated with #.

Pretreatment with the mu opioid antagonist naloxone increased oxycodone self-administration (**Figure 3B**). The ANOVA confirmed significant effects of Treatment [F(3,23)=4.128; *p*<0.05], of Time [F(3,69)=13.35; *p*<0.0001], and of the interaction [F(9,69)=2.9; *p*<0.01] and the post hoc analysis further confirmed that significantly more infusions were obtained after 0.03 mg/kg naloxone, i.p., compared with all other treatment conditions from 30-60 minutes and compared with the 0.3 mg/kg pretreatment at 90 minutes.

### 3.4 THC (i.p.) alters the self-administration of different opioid doses in parallel

In the 4 h oxycodone self-administration study, dose-related effects of THC (by i.p. injection) were observed in both adolescent exposure groups with either per-infusion dose of oxycodone (**Figure 4A, B**). There were no effects of adolescent treatment group confirmed by an initial analysis at either per-infusion dose and, collapsed across group, oxycodone self-administration was significantly affected by Oxycodone Dose [F(1,18)=21.93; *p*<0.0005] and THC Dose [F(4,72)=30.03; *p*<0.0001]. The post hoc test confirmed that significantly fewer infusions were obtained after THC injection compared with vehicle for the lower (5-10 mg/kg THC), higher (7.5-10 mg/kg THC) or collapsed across (5-10 mg/kg THC) oxycodone doses. The follow up ANOVAs also confirmed significant effects of Oxycodone dose and THC pre-treatment condition, but not of the interaction of factors, for both THC [Oxy: F(1,8)=15.92; *p*<0.005; THC: F(4,32)=15.05; *p*<0.0001] and PG [Oxy: F(1,9)=7.22; *p*<0.05; THC F(4,36)=14.68; *p*<0.0001] adolescent treatment subgroups. The post hoc test confirmed that rats in each group self-administered significantly more infusions of 0.06 mg/kg oxycodone than of 0.15 mg/kg oxycodone with pre-treatment conditions from 0.0-7.5 mg/kg, THC, i.p.. A post hoc test also confirmed that reductions in self-administration of either oxycodone dose were produced in each group by pre-session injection of THC in a dose-dependent manner. Finally, analysis of the pattern of self-administration within-session demonstrated a preserved loading dose / maintenance dose phenomenon in each of the THC pre-treatment conditions for high and low per-infusion dose tests (See **Supplementary Information; Figure S2**). A final experiment was conducted with two different per-infusions doses of heroin (0.006, 0.06 mg/kg/infusion) available under a Progressive Ratio response contingency (**Figure 4C**). The analysis confirmed a significant effect of THC pre-treatment dose [F(3,90)=8.77; *p*<0.0001] (but not of adolescent treatment group or per-infusion dose of heroin) and the Dunnett’s post hoc test further confirmed that, collapsed across per-infusion dose of heroin, significantly higher breakpoints were reached after 1.0 mg/kg THC and lower breakpoints after 10.0 mg/kg THC compared with vehicle.

**Figure 4.**
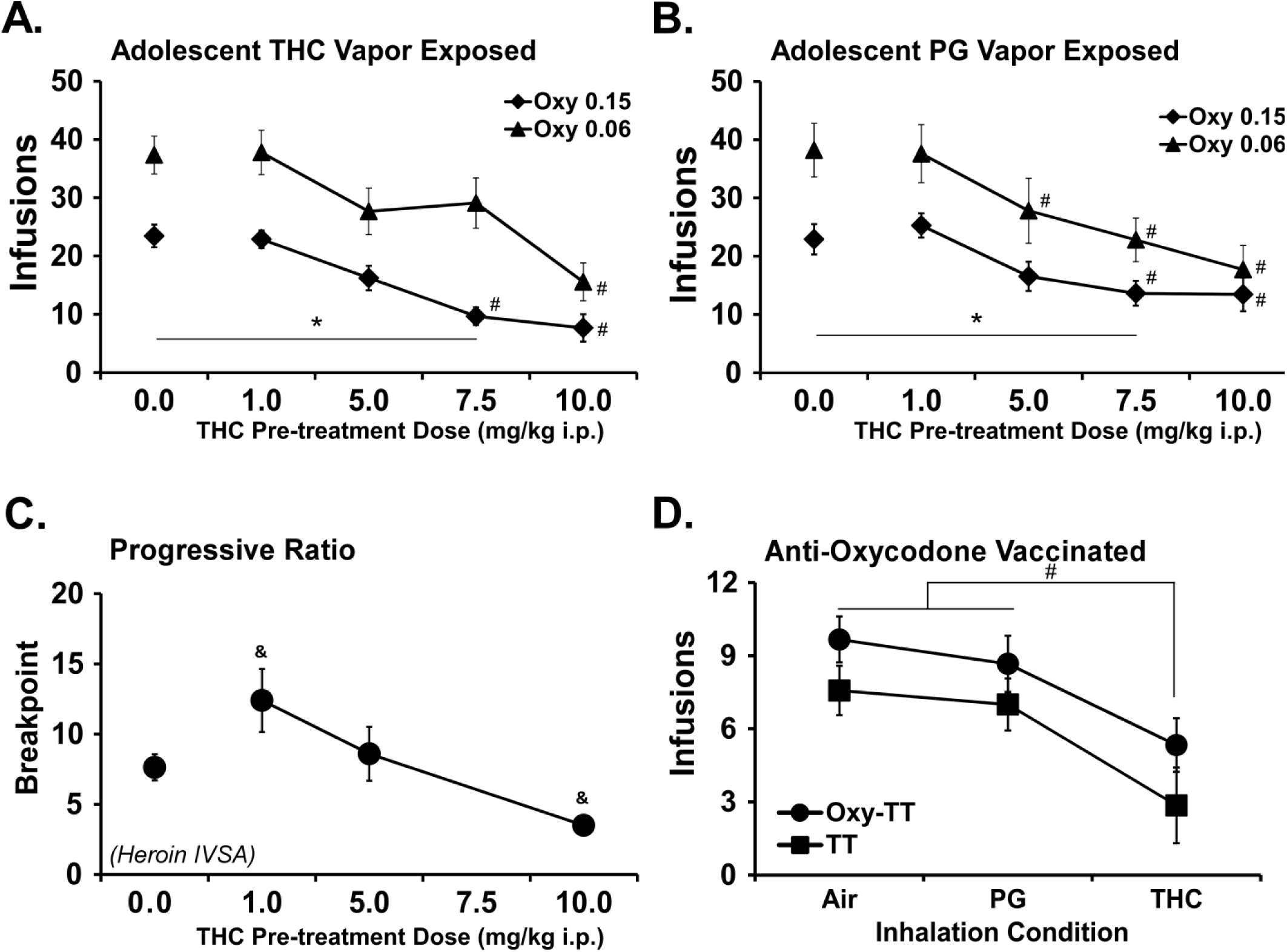
THC vapor inhalation reduces opioid self-administration under short access conditions. Mean (N=19 ±SEM) infusions of oxycodone obtained by two groups of adult rats exposed during adolescence to repeated A) THC vapor or B) PG vapor. A significant difference between the oxycodone per-infusion doses, within THC pre-treatment condition, is indicated with *. A significant difference from vehicle pre-treatment, within per-infusion dose, is indicated with #. C) Mean (N=15; ±SEM) breakpoints reached by rats permitted to self-administer heroin under a Progressive Ratio response contingency. A difference from the vehicle pre-treatment condition is indicated with &. D) Mean (TT, N=7; Oxy-TT, N=9; ±SEM) oxycodone infusions obtained following 30 minutes inhalation of Air, Propylene Glycol (PG) vehicle vapor or THC vapor. Significant differences from Air and PG vehicle condition are indicated by #.

### 3.5 THC vapor attenuates oxycodone self-administration under short access (1 h) conditions

The groups of rats that had been vaccinated with the anti-oxycodone vaccine (Oxy-TT) or the carrier protein control (TT) were trained to self-administer oxycodone in 1 h limited access sessions (see **Supplementary Information; Figure S1**). Exposure to THC vapor inhalation for 30 minutes prior to the session significantly reduced (**Figure 4D**) the number of infusions of oxycodone obtained (Significant main effect of Dose condition [F(2,28)=21.96; *p*<0.0001] but not of Group or the interaction of factors) relative to Air or PG inhalation (which did not differ from each other).

### 3.6 THC enhances oxycodone-induced antinociception

A study was next conducted to determine if injection of a combination of oxycodone and THC would produce interactive effects on antinociception (**Figure 5A**). A group of male (N=6) and female (N=8) Wistar rats were injected with the cannabinoid vehicle or THC (10 mg/kg, i.p.) 30 min prior to the saline vehicle or oxycodone (0.0 or 1 mg/kg, s.c.). The ANOVA confirmed significant effects of Time [F(3,39)=6.0; *p*<0.005], of Drug Condition [F(3,39)=41.47; *p*<0.0001], and of the interaction [F(9,117)=4.71; *p*<0.0001]. Post hoc analysis confirmed that when THC and oxycodone were administered in combination, this induced significantly longer tail withdrawal latency compared to other drug conditions at the 30 and 60 min time points. Similar effects were confirmed within the male (Time [F(3,15)=6.24; *p*<0.01]; Drug Condition [F(3,15)=44.42; *p*<0.0001]; Interaction [F(9,45)=2.76; *p*<0.05]; Post hoc: Combination > all other conditions 30-60 minutes post-injection) and female (Time [n.s.]; Drug Condition [F(3,21)=16.01; *p*<0.0001]; Interaction [F(9,63)=3.02; *p*<0.005]; Post hoc: Combination > all other conditions 30-90 minutes post-injection) subgroups.

**Figure 5.**
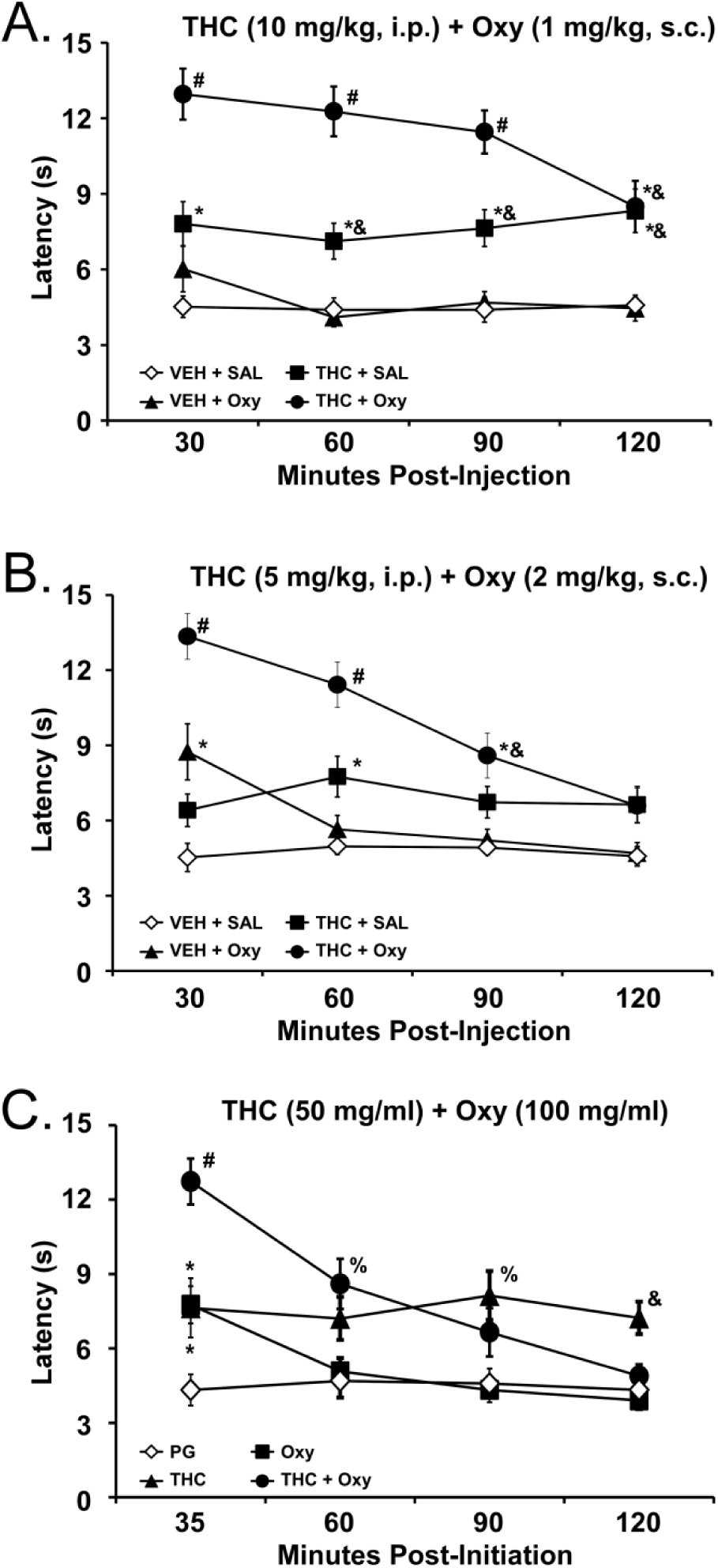
THC and oxycodone co-administration produces additive effects on antinociception. A) Mean (N=14, 8F; ±SEM) tail withdrawal latency following administration of THC (10 mg/kg, i.p.), oxycodone (1 mg/kg, s.c.) or the combination. B) Mean (N=14, 8F; ±SEM) tail withdrawal latency following administration of THC (5 mg/kg, i.p.), oxycodone (2 mg/kg, s.c.) or the combination. C) Mean (N=10; ±SEM) tail withdrawal latency following inhalation of vapor from PG, THC (50 mg/mL), Oxycodone (100 mg/mL) or the THC/Oxycodone combination. Significant difference from all other treatments is indicated with #, a significant difference from VEH+Sal or PG with *, and a significant difference from VEH+Oxy or Oxy alone with *. A significant difference from PG and Oxy alone is indicated with %.PG=propylene glycol vehicle, VEH= 1:1:8 vehicle used for THC injections; Sal=Saline vehicle used for oxycodone injections.

The rats were next injected with the vehicle or THC (5 mg/kg, i.p.) 30 min prior to the saline vehicle or oxycodone (0.0 or 2 mg/kg, s.c.) again in a randomized order (**Figure 5B**). The ANOVA confirmed significant effects of Time [F(3,156)=20.55; *p*<0.0001], of Drug Condition [F(3,52)=17.93; *p*<0.0001], and of the interaction [F(9,156)=9.21; *p*<0.0001].

Post hoc analysis confirmed that oxycodone administered alone significantly increased tail withdrawal latency (30 min), whereas the THC and oxycodone combination significantly increased latency compared to vehicle for up to 90 min. The combination of THC and oxycodone produced significantly longer withdrawal latency compared with the latency following THC (30 and 60 minutes) and oxycodone (30-90 minutes) injections alone. Similar effects of THC and oxycodone co-administration were confirmed within the male and female subgroups (see **Supplementary Information; Figure S4)**.

A separate group of rats (N=10) was tested for the antinociceptive effects of vaporized oxycodone and THC. The tail withdrawal latency was lengthened following inhalation of vaporized oxycodone (100 mg/ml), THC (50 mg/ml) or the THC:oxycodone combination (**Figure 5C**). The ANOVA confirmed significant main effects of Time after vapor initiation [F(3,27)=11.31; *p*<0.0001], of Drug Condition [F(3,27)=20.78; *p*<0.0001] and of the interaction of factors [F(9,81)=7.55; *p*<0.0001]. The post hoc analysis confirmed that inhalation of combined oxycodone and THC significantly increased tail withdrawal latency compared to PG (35-60 minutes after vapor initiation), oxycodone alone (35-60 minutes after vapor initiation) or THC alone (35 minutes after vapor initiation). Significantly increased latency compared with PG inhalation was also observed after inhalation of oxycodone alone (35 minutes after vapor initiation) or THC alone (35, 90 minutes after vapor initiation).

### 3.7 Plasma THC levels after vapor inhalation or i.p. injection

Plasma THC levels were similar following vapor inhalation of THC (100 mg/mL) for 30 minutes and injection of THC (10 mg/kg, i.p.) as depicted in **Figure 6**. Analysis was between-groups for both factors due to missing blood samples for one animal after vapor inhalation (240 minutes after the start of inhalation) and i.p. injection (120-240 minutes post-injection); a significant main effect of Time was confirmed [F(3,37)=23.28; *p*<0.0001]. (A preliminary repeated measures analysis with the N=5 with complete data produced no difference in significant effects.) The Tukey post hoc test confirmed that plasma THC levels were significantly lower than the 35 minute timepoint after i.p. injection (120-240 minutes) or vapor inhalation (60-240 minutes). Plasma THC was also significantly lower 240 minutes after injection compared with the 60 minute time-point.

**Figure 6:**
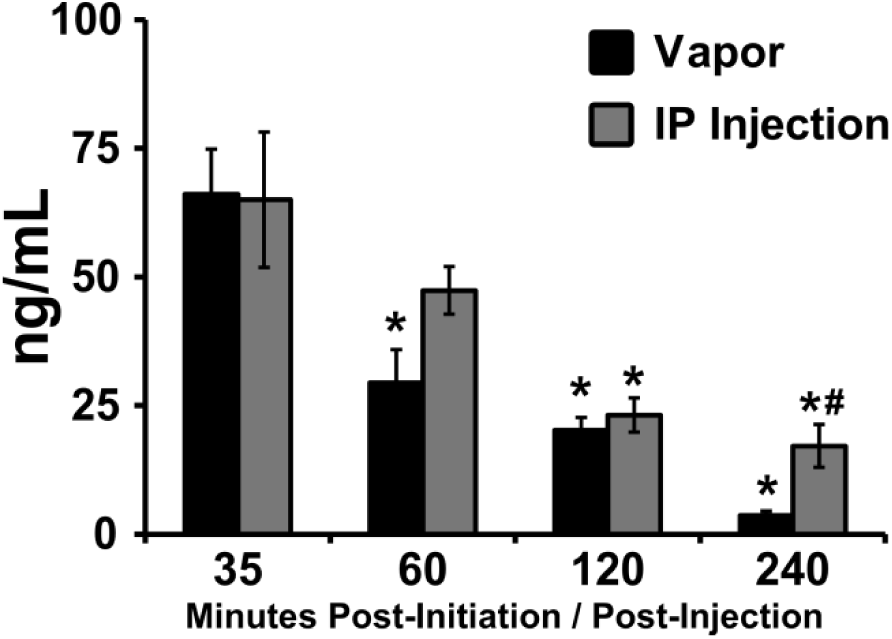
Plasma THC concentration following vapor inhalation or injection. Mean (N=5-6; ±SEM) plasma THC following injection (10 mg/kg, i.p.) and vapor inhalation (THC 100 mg/mL for 30 minutes). A significant difference from the 35 minute time-point is indicated by * and a difference from the 60 minute time-point with #.

## 4. Discussion

This study found that THC reduced the number of infusions of oxycodone that were self-administered under either short or extended daily access conditions, and increased in the magnitude and duration of antinociception produced by oxycodone. The data support a conclusion that THC interacts with the effects of oxycodone when the two drugs are co-administered such that THC enhances the effects of a given dose of oxycodone, *in vivo*. It is particularly notable that THC reduced the self-administration of oxycodone even in rats trained to escalated levels using an extended access (8 h) paradigm. The effects on opioid reward were lasting, since a single THC delivery by injection or inhalation 30 min prior to the sessions significantly reduced the self-administration of oxycodone for up to 5 hours. THC-mediated enhancement of the antinociceptive effects of oxycodone was likewise present across both routes of administration. The impact of THC was dose-dependent and it generalized across the injection and inhalation routes of administration under experimental conditions which produced similar plasma THC levels (**Figure 6**).

Additional studies demonstrated that the effects of THC on oxycodone self-administration were specific, and not merely due to behavioral sedation, by using two strategies which reduced the amount of oxycodone reaching the brain on each self-administered infusion. In both cases, THC produced a parallel shift in self-administered oxycodone. First, we used an anti-oxycodone vaccine strategy which results in about a 50% reduction in brain levels of oxycodone after a given dose (Nguyen et al., 2018c) and increases the intravenous self-administration of oxycodone under easy access conditions (i.e., a fixed-ratio 1 response contingency), albeit slightly less than required to compensate for reduced brain levels. THC vapor inhalation reduced self-administration similarly in each vaccine group. An additional study used reduction of the available per-infusion dose of oxycodone, and a broader range of injected THC doses, to further illustrate the point. The essentially parallel dose-response functions for i.p. THC pre-treatment under the 0.06 and 0.15 mg/kg oxycodone maintenance conditions (**Figure 4 A,B**) is incompatible with a simple interpretation of a sedative effect of THC and further supports the conclusion that the THC enhanced the reinforcing efficacy of a given dose of oxycodone. Relatedly, the bin analysis for the 4 h groups (see **Supplementary Information; Figure S2**) demonstrated preservation of a loading / maintenance dose pattern. Furthermore, most individuals obtained at least one infusion of oxycodone (0.06 mg/kg/inf) in the first 15 minutes of the session after 5 (17/19 rats) or 7.5 (16/19) mg/kg THC; these were doses which produced significant reductions in self-administration across the group. Finally, we showed that a low dose of THC that does not affect opioid self-administration under a FR response contingency increases breakpoints when animals self-administer heroin under a PR contingency (**Figure 4C**). This pattern is interpreted as enhanced reward and the increased behavioral output is also incompatible with interpretations of behavioral suppression. Relatedly, neither injected (10 mg/kg, i.p.) nor inhaled (1-200 mg/mL, 30 minutes) THC consistently reduces spontaneous locomotor activity in our hands (Nguyen et al., 2016b; Taffe et al., 2015); also see (Wiley and Burston, 2014; Wiley et al., 2007). The Craft lab has reported some locomotor suppressive effects of 10 mg/kg THC in rats (Britch et al., 2017; Craft and Leitl, 2008), however those studies were conducted in the rats’ inactive (light) cycle of the day unlike the present studies which were conducted in the active (dark) cycle. Therefore the most parsimonious conclusion is that in this study THC enhanced the rewarding value of opioids, as opposed to altering behavior in a non-specific manner.

As a minor caveat, this conclusion contrasts with a study in which THC and the CB1 full agonist CP55940 produced rate-suppressant effects when given in combination with remifentanil, but did not appear to enhance the reinforcing effects, in rhesus monkeys responding under a food/drug choice procedure (Maguire and France, 2018). Future follow up would be required to determine if the difference is attributable to the IVSA drug identity (i.e. heroin vs. oxycodone), experimental species, the route and chronicity of THC exposure, both or some other factor. Similarly, the lack of lasting effect of adolescent THC exposure on oxycodone self-administration contrasts with a study in which THC injection every third day from PND 28-49 increased the IVSA of heroin, initiated on PND 57 (Ellgren et al., 2007) but agrees with another study which found no effect of adolescent THC on heroin IVSA acquisition (Stopponi et al., 2014). For the present study, however, the key result is that THC was equally effective at reducing the IVSA of oxycodone in each of the adolescent-treatment groups, suggesting no lasting alteration of CB1 function (Nguyen et al., 2018a; Nguyen et al., 2018b) that was relevant to the interaction of THC with the rewarding effects of oxycodone.

Analysis of the within-session patterns of responding in the 4 h study found that significant differences in oxycodone self-admin lasted up to 2 h in both high and low per-infusion doses. More importantly the assessment of the self-administration pattern of oxycodone in the first four 15 min bins demonstrated a loading dose phenomenon in each of the THC pre-treatment conditions for high and low per-infusion dose tests (**Figure S2**). The consistency of this pattern across the THC pre-treatments is further evidence that effects of THC were not merely sedative or cataleptic. Relatedly, we have previously shown that 10 mg/kg, THC, i.p. does not suppress male rat activity for up to 4 hours after injection and that a 20 mg/kg dose does not suppress activity until 3 h post-injection (Taffe et al., 2015). Furthermore, while the inhalation of THC vapor for 30 minutes lowers activity rate in some, but not all cases, it fails to make rats completely immobile (Javadi-Paydar et al., 2018; Nguyen et al., 2016b).

In addition to the dose-dependency of the effect of THC on oxycodone IVSA, an interpretation of mechanistic specificity is further enhanced by the finding that prior administration of the CB1 antagonist SR 141716 blocked the effects of THC inhalation. These data can be considered with the finding that administration of the mu opioid receptor agonist oxycodone decreased, whereas the antagonist naloxone increased, the amount of oxycodone that was self-administered. This supports the conclusion that the mechanism by which THC decreases oxycodone IVSA is mediated by the CB1 receptor and produces an enhancement of the effects of oxycodone at the mu opioid receptor. Opposing the effects of oxycodone would increase, rather than decrease, self-administration as was found after naloxone pre-treatment.

The study also showed that sensation for a noxious stimulus, as classic preclinical model of analgesic activity, was additively diminished by the co-administration of THC with oxycodone compared with either drug administered alone. Prior work has shown antinociceptive interactions between mu-opioid and cannabinoid receptor ligands in formalin test of inflammatory pain (Yuill et al., 2017) and in nociception rhesus monkeys (Li et al., 2008). The present study extends those results to the interaction of THC with the effects of the prescription opioid oxycodone. It was notable that the interactive antinociceptive effects of THC with oxycodone appeared to last long past the duration of activity of oxycodone administered by itself, which was only about 30 minutes. This may suggest a second benefit of adding THC to oxycodone (i.e., extended duration of action) in addition to the primary effect, (i.e., a reduction of dose to produce comparable immediate effect).

In conclusion, this study showed for the first time that THC interacts with oxycodone within preclinical models of both reward and analgesia. The totality of the results presented attribute this most parsimoniously to an additive effect whereby THC enhances the effectiveness of a given dose of oxycodone. This provides experimental evidence for the likely pharmacological specificity of epidemiological findings, i.e. from medical marijuana states. The evidence that THC can attenuate the self-administration of oxycodone suggests a potential therapeutic effect akin to agonist therapy for those attempting to reduce nonmedical oxycodone use. The nociception data also suggest that co-use of cannabis and prescriptions opioids such as oxycodone might provide effective pain control with lower doses than would be required for either drug alone. One critical next step might be to determine the extent to which oxycodone self-administration in a chronic pain state could be modified by THC. Overall, this study shows that further investigation of cannabinoid / opioid interactions may identify improved therapeutic approaches for analgesia and possible mechanisms to reduce opioid addiction.

## Acknowledgements

The authors are grateful to Tony M. Kerr for the plasma analysis and to Shawn M. Aarde for contributions to the invention and initial validation of the vapor inhalation method. This is manuscript #29627 from The Scripps Research Institute.

