## Supplementary Materials for "Δ^9^-tetrahydrocannabinol Attenuates Oxycodone Self-Administration Under Extended Access Conditions"

Running Title: THC / Oxycodone interactions

### Supplemental Methods

**Intravenous Catheterization.** Rats were anesthetized with an isoflurane/oxygen vapor mixture (isoflurane 5% induction, 1-3% maintenance) and prepared with chronic indwelling intravenous catheters as described previously (Nguyen *et al*, 2017c). The intravenous catheters consisted of a 14.5-cm length of polyurethane based tubing (Micro-Renathane®, Braintree Scientific, Inc, Braintree, MA) fitted to a guide cannula (Plastics One, Roanoke, VA) curved at an angle and encased in dental cement anchored to an ~3 cm circle of durable mesh. Catheter tubing was passed subcutaneously from the animal's back to the right jugular vein. Catheter tubing was inserted into the vein and tied gently with suture thread. A liquid tissue adhesive was used to close the incisions (3M™ Vetbond™ Tissue Adhesive: 1469SB, 3M, St. Paul, MN). A minimum of 4 days was allowed for surgical recovery prior to starting an experiment. For the first three days of the recovery period, an antibiotic (cefazolin) and an analgesic (flunixin) were administered daily. During testing and training, intravenous catheters were flushed with ~0.2-0.3 ml heparinized (166.7 USP/ml) saline before sessions and ~0.2-0.3 ml heparinized saline containing cefazolin (100 mg/mL) after sessions.

Catheter patency was assessed once a week after the last session of the week, via administration through the catheter of ~0.2 ml (10 mg/ml) of the ultra-short-acting barbiturate anesthetic Brevital sodium (1% methohexital sodium; Eli Lilly, Indianapolis, IN). Animals with patent catheters exhibit prominent signs of anesthesia (pronounced loss of muscle tone) within 3 sec after infusion. Animals that failed to display these signs were considered to have faulty catheters, and if catheter patency failure was detected, data that were collected after the previous passing of this test were excluded from analysis.

**Inhalation Apparatus and Procedure:** Sealed exposure chambers were modified from the 254mm X 267mm X 381mm Allentown, Inc (Allentown, NJ) rat cage to regulate airflow and the delivery of vaporized drug to rats, as has been previously described (Nguyen *et al*, 2016a; Nguyen *et al*, 2016b). An e-vape controller (Model SSV-1; La Jolla Alcohol Research, Inc, La Jolla, CA, USA) was triggered to deliver the scheduled series of puffs (4 10 sec puffs every 5 minutes) from Protank 3 Atomizer (Kanger Tech; Shenzhen Kanger Technology Co.,LTD; Fuyong Town, Shenzhen, China) e-cigarette cartridges for the 8 h self-administration experiment. Type 2 sealed exposure chambers (La Jolla Alcohol Research, Inc; La Jolla, CA, USA) and a second generation e-vape controller (Model SSV-2; La Jolla Alcohol Research, Inc, La Jolla, CA, USA) with Herakles Sub Ohm Tank e-cigarette cartridges (Sense; Shenzhen Sense Technology Co., LTD; Baoan Dist, Shenzhen, Guangdong, China) controlled by MedPC IV software (Med Associates, St. Albans, VT USA) were used to deliver vapor (1 10 second puff every 5 minutes) for tail withdrawal and the 1 h self-administration experiments. The chamber air was vacuum controlled by a chamber exhaust valve (i.e., a “pull” system) to flow room ambient air through an intake valve at ~1 L per minute. This also functioned to ensure that vapor entered the chamber on each device triggering event. The vapor stream was integrated with the ambient air stream once triggered. For self-administration studies, rats were exposed to 30 min of THC vapor inhalation (followed by a 5 min period for chamber clearance) immediately prior to the start of self-administration sessions.

**Hapten Synthesis, Vaccine Formulation and Administration.** The oxycodone hapten (Oxy) was designed with an activated linker extending from the bridgehead nitrogen to directly react with the surface lysines of carrier protein tetanus toxoid (TT) or BSA. Oxycodone hapten was synthesized according to previously published methods from the Janda laboratory with slight modification in the reductive amination and amide bond formation steps (Kimishima *et al*, 2016; Nguyen *et al*, 2017d; Nguyen *et al*, 2018). Vaccines were formulated the day of immunization using 13:1:5 (v/v/v) mixture of Oxy-TT (1.0 mg/ml in PBS) or control TT (1.0 mg/ml in PBS), CpG ODN 1826 (5 mg/ml in PBS), and Alhydrogel® (alum, 10 mg/ml, InvivoGen) and administered intraperitoneally. Rats were administered the conjugate vaccine (Oxy-TT; N=12) or tetanus toxoid only (TT; N=10) on Weeks 0, 2, and 4, adapted from a vaccination protocol previously reported (Nguyen *et al*, 2016c; Nguyen *et al*, 2017a). Rats were prepared with chronic intravenous catheters on week 7 of the vaccination protocol and allowed one week of recovery prior to the start of self-administration experiments. Within the TT group, 8 rats completed acquisition with patent catheters and 7 completed the THC inhalation study. Within the Oxy-TT group, 11 completed the entire study.

**Adolescent Vapor Exposure.** The adolescent rats were divided into two groups of 12 which received 30 minute episodes of vapor exposure, b.i.d., qa. 5 h, in pairs to either THC (100 mg/mL) or the propylene glycol (PG) vehicle on sequential days PND 35-39 and again on PND 42-46. These groups received intravenous catheter implant surgery on PND 84-87 and initiated IVSA of oxycodone (0.15 mg/kg/infusion; 8 h sessions; Fixed Ratio 1 response contingency) on PND 112. Following an acquisition interval of 17 sessions the rats completed six sessions of Fixed Ratio (8 h) and Progressive Ratio (3 h) dose substitution (0.006, 0.06, 0.15 mg/kg/infusion) in a counter-balanced order. Thereafter the rats were returned to 4 h sessions of oxycodone IVSA (0.15 mg/kg/infusion; FR1) for the current study.

**Plasma THC analysis.** Blood samples were collected ( ~500 µl) from the indwelling catheters. Plasma THC content was quantified using fast liquid chromatography/mass spectrometry (LC/MS) adapted from (Irimia *et al*, 2015; Lacroix and Saussereau, 2012; Nguyen *et al*, 2017b). 50 µl of plasma were mixed with 50 µl of deuterated internal standard (100 ng/ml CBD-d3 and THC-d3; Cerilliant), and cannabinoids were extracted into 300 uL acetonitrile and 600 µl of chloroform and then dried. Samples were reconstituted in 100 µl of an acetonitrile/methanol/water (2:1:1). Separation was performed on an Agilent LC1100 using an Eclipse XDB-C18 column (3.5um, 2.1mm x 100mm) using gradient elution with water and methanol, both with 0.2 % formic acid (300 µl/min; 73-90%). THC was quantified using an Agilent MSD6140 single quadrodpole using electrospray ionization and selected ion monitoring [THC (m/z=315.2) and THC-d3 (m/z=318.2)]. Calibration curves were conducted daily for each assay at a concentration range of 0-200 ng/mL and observed correlation coefficients were 0.999.

### Supplemental Results

#### Acquisition of oxycodone self-administration in vaccinated groups

The TT (N=8) and Oxy-TT (N=11) vaccinated rats were trained to self-administer oxycodone across 15 session with group differences observed only under the FR1 response contingency (**Figure S1**). Oxy-TT group self-administered more oxycodone during the FR1 phase of the acquisition, consistent with a sequestration of part of the dose in the bloodstream. The ANOVA confirmed significant effects of Session [$F(14,238)=41.13$; $p<0.0001$], of vaccine Group [$F(1,17)=6.23$; $p<0.05$] and of the interaction of Group with Session [$F(14,238)=4.21$; $p<0.0001$] on oxycodone intake. The post hoc test confirmed that the Oxy-TT group obtained more infusions for sessions 9, 11, 12, 14.

The Oxy-TT vaccinated animals self-administered more oxycodone than the TT controls under FR1 response contingency conditions and about the same number of infusions under a PR contingency following the acquisition period. This is consistent with two similarly vaccinated groups in a prior finding (Nguyen *et al*, 2018) and is likely a behavioral marker of the ~50% decrease in brain oxycodone that is produced. In the present study, the relative impact of THC inhalation to suppressed IVSA was similar in each group and, if anything, slightly lesser in the Oxy-TT group. This outcome is also consistent with an effect of THC on the rewarding value of self-administered oxycodone rather than a general behavioral suppression.

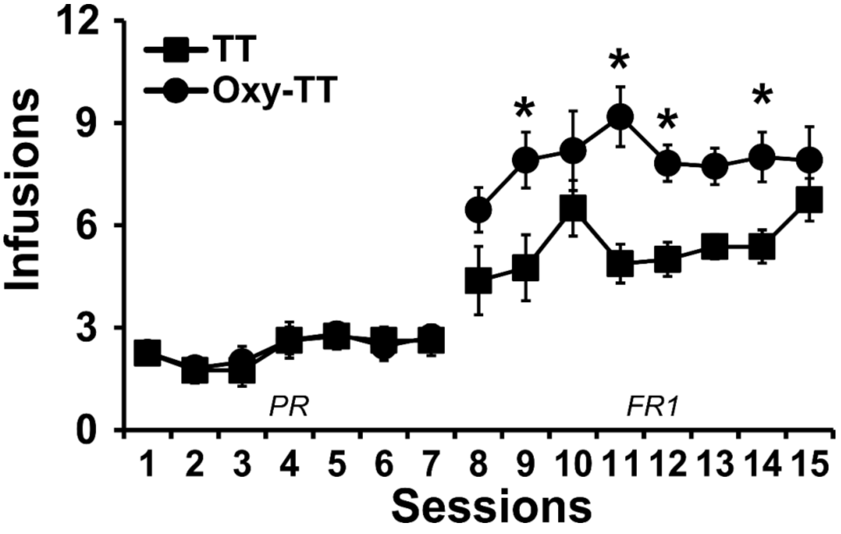

**Figure S1.** *A) Mean infusions obtained by groups of male rats vaccinated with the tetanus toxioid carrier protein (TT, N=8; ±SEM) or the anti-oxycodone conjugate vaccine (Oxy-TT, N=11; ±SEM) or trained to self-administer oxycodone (0.15 mg/kg/inf) within 1 h sessions under a Progressive Ratio (PR; Sessions 1-7) or Fixed Ratio 1 (FR1; Sessions 8-15) response contingency. Significant differences within group from session 1 are indicated by *. Significant differences from Air and PG vehicle condition are indicated by #.*

***THC dose-dependently reduces initial loading phase of oxycodone self-administration***

In the repeated THC vapor study, further analysis was conducted collapsed across adolescent treatment group because there were no significant differences associated with group confirmed in an analysis that included this as a factor (**Figure S2**). Significant effects of THC pre-treatment, hour bin and the interaction of factors were confirmed for the 0.06 mg/kg [THC: $F(4,72)=17.67$; $p<0.0001$; Hour Bin: $F(3,54)=11.46$; $p<0.0001$; Interaction: $F(12,216)=3.39$; $p<0.0005$] and 0.15 mg/kg [THC: $F(4,76)=21.72$; $p<0.0001$; Hour Bin: $F(3,57)=19.05$; $p<0.0001$; Interaction: $F(12,228)=2.91$; $P<0.001$] conditions. The

**Figure S2.** *A,B) Mean (N=19; ±SEM) obtained by rats self-administering 0.06 or 0.15 mg/kg/inf oxycodone following THC injection (0-10 mg/kg, i.p.) during 4 hour sessions. C,D) Analysis of the loading phase (first hour, by 15 minute bins) showed dose-dependent reductions in oxycodone infusions. A significant difference from the first time-point within pre-treatment condition is indicated with &, the bar indicates this is true for all treatment conditions. Significant differences from the vehicle are indicated with *, from the vehicle and 1.0 mg/kg dose with #, and differences from all other doses with $.*

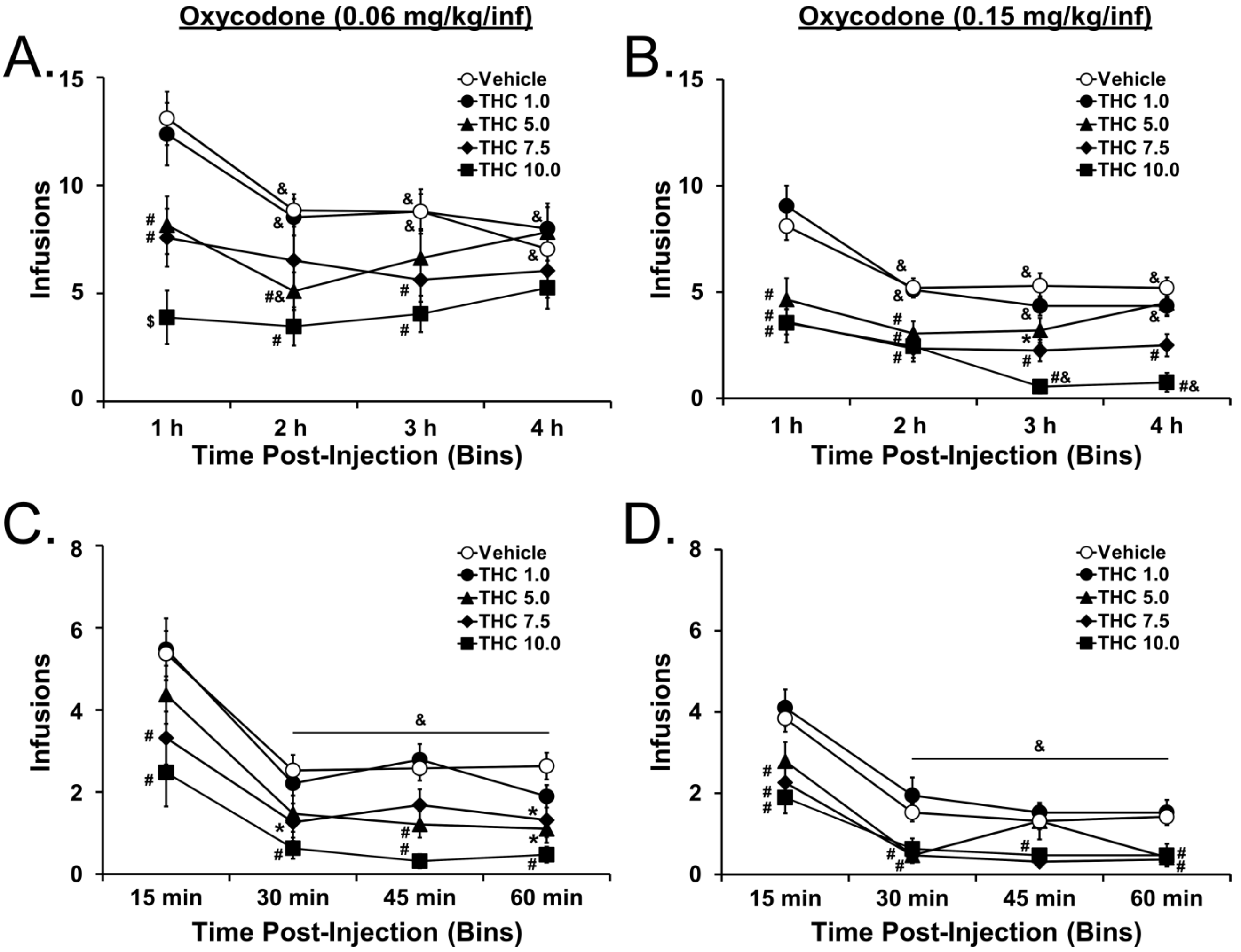

post-hoc test confirmed that intake in the first two hours differed across treatment conditions. This showed that significant differences in oxycodone self-admin lasted 2 h in both high and low per-infusion doses. Assessment of the first hour by 15 min bins demonstrated a loading dose phenomenon in each of the THC pre-treatment conditions for high and low per-infusion dose tests. Significant effects of THC pre-treatment and 15 min bin were confirmed for the 0.06 mg/kg [THC: $F(4,72)=16.04$; $p<0.0001$; 15 min Bin: $F(3,54)=33.21$; $p<0.0001$] and 0.15 mg/kg [THC: $F(4,76)=11.55$; $p<0.0001$; 15 min Bin: $F(3,57)=75.6$; $p<0.0001$] conditions. The post-hoc tests confirmed that significantly more infusions were obtained in the first 15 minutes, compared with all three subsequent time bins, across THC pre-treatment dose conditions for each of the oxycodone doses.

***THC also alters heroin self-administration***

In a follow up study these individuals were switched to IVSA of heroin (0.06 mg/kg/inf; FR1) in 2 h sessions for four sessions and then subjected to a dose substitution study in which doses (0.0, 0.006, 0.06 and 0.15) were evaluated in a counterbalanced order. Following this rats were switched to a training dose of 0.006 mg/kg heroin and pre-treated (30 min before sessions) with doses of THC (0.0, 1.0, 7.5, 10.0 mg/kg, i.p.) in a counterbalanced order (**Figure S3A**). The ANOVA confirmed a significant effect of THC pre-treatment dose [$F(3,45)=15.95$; $p<0.0001$] and the Tukey post-hoc confirmed that this was attributable to fewer infusions of heroin being obtained after the higher two doses as depicted in **Figure S3A**. Next, rats were evaluated with THC (5.0 mg/kg, i.p.) or vehicle pre-treatment with available heroin doses of 0.006 or 0.06 mg/kg/infusion under an FR1 contingency (**Figure S3B**). The initial three-factor ANOVA failed to confirm any effect of adolescent treatment group so the main analysis collapsed across this factor. The ANOVA confirmed significant effects of per-infusion dose [$F(1,15)=17.57$; $p=0.0008$] and THC pre-treatment condition [$F(1,15)=6.05$; $p=0.0265$]. Finally, rats were evaluated in a the PR contingency procedure (3 h sessions) with THC (0.0, 1.0, 5.0, 10.0 mg/kg, i.p..) pre-treatment with available heroin doses of 0.006 or 0.06 mg/kg/infusion in a counter balanced order (**Figure S3C**). The analysis confirmed a significant effect of dose $F(3,90)=8.77$; $p<0.0001$ and the Dunnett's post hoc test further confirmed that, collapsed across per-infusion dose of heroin, significantly higher breakpoints were reached after 1.0 mg/kg THC and lower breakpoints after 10.0 mg/kg THC compared with vehicle.

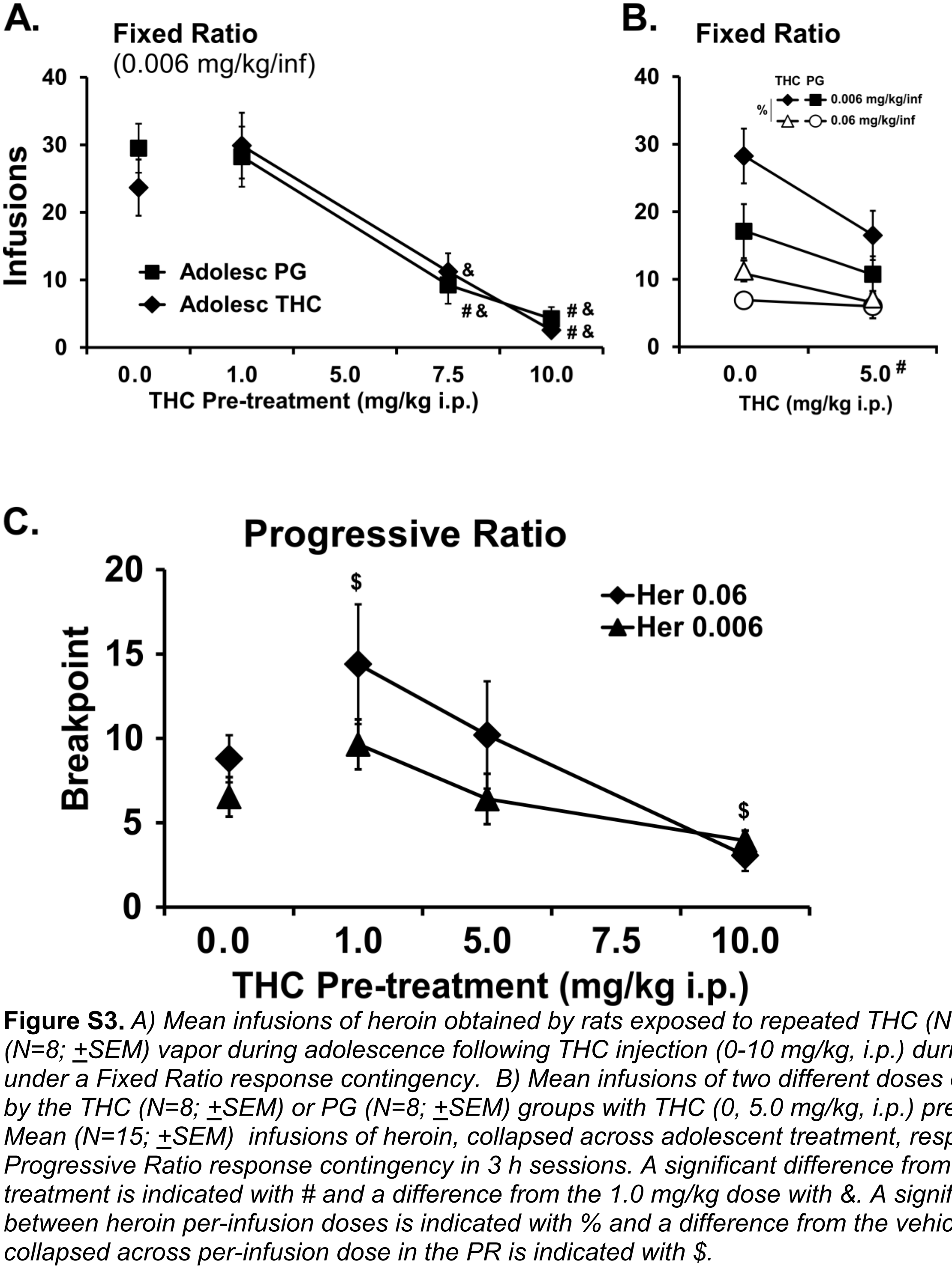

**Figure S3.** *A) Mean infusions of heroin obtained by rats exposed to repeated THC (N=9; ±SEM) or PG (N=8; ±SEM) vapor during adolescence following THC injection (0-10 mg/kg, i.p.) during 2 hour sessions under a Fixed Ratio response contingency. B) Mean infusions of two different doses of heroin obtained by the THC (N=8; ±SEM) or PG (N=8; ±SEM) groups with THC (0, 5.0 mg/kg, i.p.) pre-treatment. C) Mean (N=15; ±SEM) infusions of heroin, collapsed across adolescent treatment, responding under Progressive Ratio response contingency in 3 h sessions. A significant difference from vehicle pre-treatment is indicated with # and a difference from the 1.0 mg/kg dose with &. A significant difference between heroin per-infusion doses is indicated with % and a difference from the vehicle pre-treatment collapsed across per-infusion dose in the PR is indicated with $.*

**THC enhances oxycodone-induced antinociception in male and female rats**

Similar effects of THC, oxycodone and the combination of drugs were confirmed within the male (Time [F(3,60)=15.66; p<0.0001]; Drug Condition [F(3,20)=13.5; p<0.0001]; Interaction [F (9, 60) = 4.7; P<0.0001]; Post hoc: Combination > all other conditions 60 minutes post-injection) and female (Time [F(3,84)=6.83; p<0.0005]; Drug Condition [F(3,28)=10.37; p<0.0001]; Interaction [F(9,84)=5.07; p<0.0001] ; Post hoc: Combination > all other conditions 30 minutes post-injection) subgroups

**Figure S4.** *A,B) Mean (Male, N=6; Female, N=8; ±SEM) tail withdrawal latency following administration of THC (10 or 5 mg/kg, i.p.), oxycodone (1 or 2 mg/kg,i.p.) or the THC:oxycodone combination.*

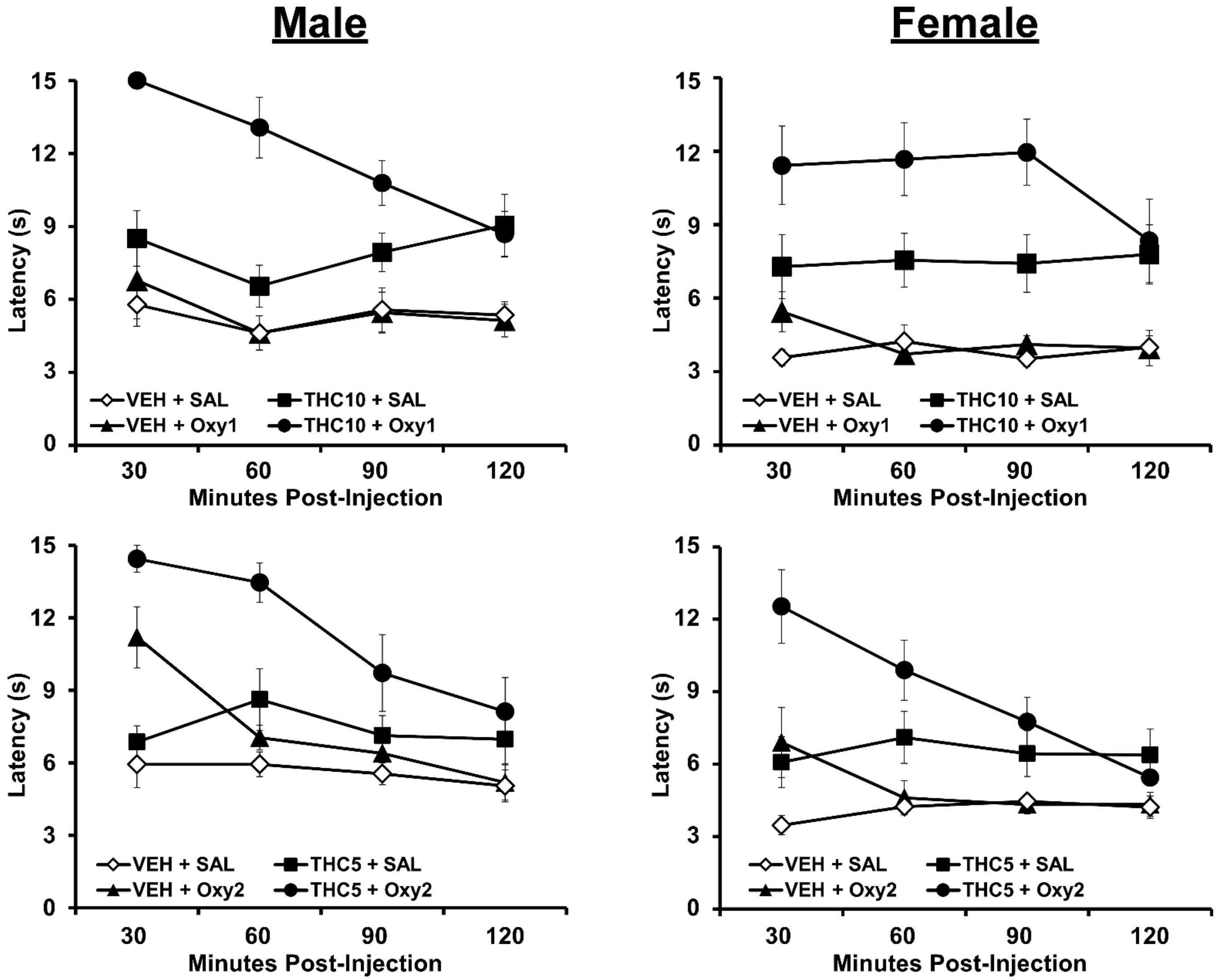
